# An ultra-sensitive and easy-to-use multiplexed single-cell proteomic analysis

**DOI:** 10.1101/2022.01.02.474723

**Authors:** Lei Gu, Ziyi Li, QinQin Wang, HuiPing Zhang, YiDi Sun, Chen Li, Hui Wang

## Abstract

Proteins analysis from an average cell population often overlooks the cellular heterogeneity of expressed effector molecules, and knowledge about the regulations of key biological processes may remain obscure. Therefore, the necessity of single-cell proteomics (SCP) technologies arises. Without microfluidic chip, expensive ultrasonic equipment, or reformed liquid chromatogram (LC) system, we established an Ultra-sensitive and Easy-to-use multiplexed Single-Cell Proteomic workflow (UE-SCP). Specifically, the flexible sorting system ensured outstanding cell activity, high accuracy, remarkable efficiency, and robustness during single-cell isolation. Multiplex isobaric labeling realized the high-throughput analysis in trapped ion mobility spectrometry coupled with quadrupole time-of-flight mass spectrometry (timsTOF MS). Using this pipeline, we achieved single-cell protein quantities to a depth of over 2,000 protein groups in two human cell lines, Hela and HEK-293T. A small batch experiment can identify and quantify more than 3200 protein groups in 32 single cells, while a large batch experiment can identify and quantify about 4000 protein groups in 96 single cells. All the 128 single cells from different cell lines could been unsupervised clustered based on their proteomes. After the integration of data quality control, data cleaning, and data analysis, we are confident that our UE-SCP platform will be easy-to-marketing popularization and will promote biological applications of single-cell proteomics.

## Introduction

Single-cell analysis characterizes each individual cell and thus has enabled the discovery and classification of previously unknown cell states.^1^ At the genomic and transcriptomic levels, single-cell sequencing has become a powerful tool for studying cell heterogeneity and identifying different phenotypic cell types. However, single-cell proteomics (SCP) based on mass-spectrometry (MS) technologies is still in its infancy due to the development of MS sensitivity and resolution, the complexity of proteome, and its non-amplifiable nature. ^1, 2^ Although antibody-based methods or fluorescent protein-based methods are widely utilized to characterize and quantify proteins at single-cell level. Antibody-based methods can generally applicable to intermediate number of proteins at once, while fluorescent protein-based methods are highly specific and facilitate monitoring protein levels over time.^3^ But both cannot provide quantifying thousands of proteins in an unbiased manner.^3, 4^ As a contrast, MS-based technologies can give comprehensive, unbiased, and discovery-oriented for the detection of proteins at single-cell level.^3-13^

Most of the existing MS-based SCP approaches are dependent on special homemade pre-processing devices, microfluidic chips, or modified liquid chromatography systems requiring special maintenance, such as the integrated single-cell analysis device (iPAD-1) developed by Xi Shao *et al*.,^12^ the nanoscale oil-gas droplet (OAD) chip developed by Zi-Yi Li *et al*.,^6^ and the nanoPOTS platform developed by Ying Zhu *et al*.^5, 14^ All those ingenuities can realize the extraction, enzymatic hydrolysis, and mass spectrometry detection of proteome at single-cell level, but special micro capillary tubes and injection valves or special microfluidic chips need to be customized in the pre-treatment process.

Carrier (Booster) channel, blank channel and single-cell channels labeled by isobaric labels (For example, tandem mass tags, TMT) can realize multiplexed single-cell proteomes collected by mass spectrometry (SCoPE-MS).^5, 7^-^10, 13^ Till now, only high-resolution MS (Orbitrap MS) have applications for this method. Not all main flow commercial mass spectrometers are suitable for the current technological process, such as timsTOF MS with higher sensitivity and better ionization efficiency. The identified and quantified protein groups in single conventional cell (10-20 μm diameter) have been reported to over 1000 using Orbitrap MS.^8, 9, 13^ Thus, MS sensitivity, resolution, and ionization efficiency are all need to be considered for upgrading and promoting this technology.

The ability to isolate adequate numbers of viable single cells is one of the key determinants in the field of single-cell omics. Among the existing SCP technologies, isolation of single-cell can be achieved through manually micro-manipulation, microfluidic devices, and fluorescence-activated cell sorting (FACS). But, the sorting throughput, accuracy or impact on living cells remains to be discussed.^15-17^ Recently, robotic automation solutions, such as acoustic microdispensing systems (cellenONE^®^, Cellenion), piezoelectric approaches (WOLF Cell Sorter, NanoCellect Biomedical or SingleCell Printer, Cytena), allow for high-throughput in single-cell isolation and have been used in single-cell genomic and transcriptomic analyses.^16-18^ Among them, the cellenONE^®^ system, integrating the microcapillary dispersion technology with constant imaging of the target cells, offers a new, innovative approach to isolate cells gently, rapidly and with high precision.^16^ Besides, this system can be applied to a wide range of cell samples with diameters up to 70 μm in practical. Meanwhile, the initial number of isolated cells is also allowed to be as low as a few dozen. Whether it is rare cells, immune cells, or cellular heterogeneity analysis, this system can be competent. Therefore, we extend the cellenONE^®^ system to the field of single-cell proteome technology in this work.

In order to reduce the application threshold, expand instrument compatibility, and improve the precision of single-cell sorting, we develop an Ultra-sensitive and Easy-to-use multiplexed Single-Cell Proteomic analysis (UE-SCP). Using the UE-SCP workflow, more than 2000 protein groups can be identified and quantified in a single HELA or HEK293T cell, which is among the best SCP methods reported so far.

## Results

The UE-SCP workflow is shown in Figure 1. The main differences from SCoPE-MS series ^7, 8,11^ are lysis buffer selection, microporous plate choice, cell sorting system, and LC-MS/MS instrument. We try to increase analytical sensitivity and lower the technological barriers through multiple ways. First, commercial microporous plates were used to collect the isolated single cells instead of homemade chips. Second, the cellenONE^®^ is used to robotic automation sort single cells from cell suspension, scatheless and efficiently. Third, timsTOF MS, with higher sensitivity and better ionization efficiency, is optimized to do TMT labelling, MS data acquisition, and database searching and data analysis in our SCP platform.

**Figure 1.**
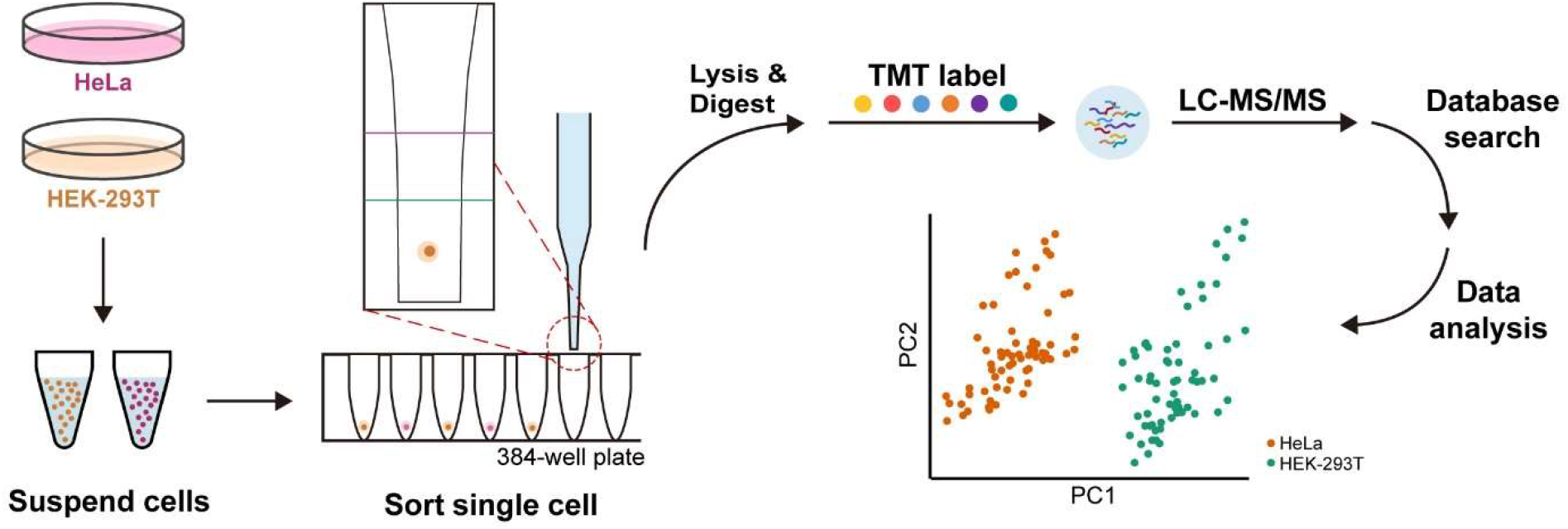
Workflow of the Ultra-sensitive and Easy-to-use multiplexed Single-Cell Proteomic analysis (UE-SCP). Single cells are sorted into 384-well plate through the cellenONE^®^, then lysed, digested, labelled, combined, and analyzed by LC-MS/MS. One TMT set contains four single-cell channels, one blank channel and one carrier channel.

### Single-cell sorting based on cellenONE^®^ system

After brief scanning, the cellenONE^®^ system can be set to dispense one single HeLa or HEK-293T cell per well in 384 well-plates using optimized isolation parameters of given cell suspension. Figure 2A and 2B show the representative real time images for isolated single HeLa and single HEK-293T cell. The cell collection scatterplots for Hela cell or HEK-293T cell suspension are drawn in Figure 2C and 2D, respectively. The cell sorting experiments present high efficiency and super accuracy. Isolation frequency is as fast as 4 Hela cells or 8 HEK-293T cells per min, and both recovery rates are round to 50% in the given cell suspensions.

**Figure 2.**
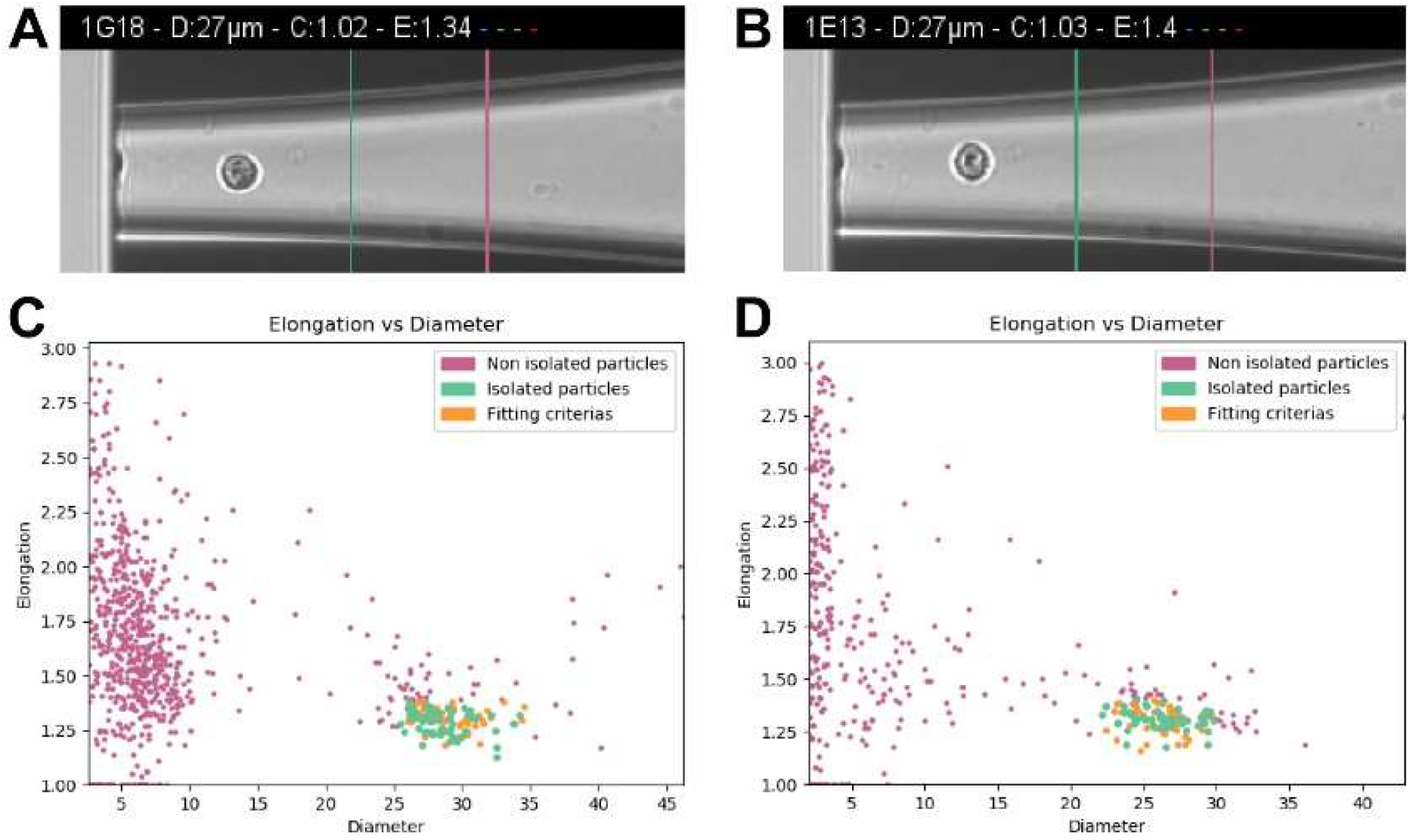
Suspend Hela or HEK-293T cells were isolated to single-cell based on optical detection. The isolated single HeLa (A) and single HEK-293T (B) cell during flexible sorting using the cellenONE^®^ system. Summarized cell collection scatterplots were shown in panel C (Hela) and panel D (HEK-293T).

### Optimizing LC-MS/MS conditions for TMT-labeled SCP pipeline using defined ratios of HEK-293T tryptic digests

As one of the mainstream mass spectrometers, timsTOF MS has higher sensitivity and better ionization efficiency. We firstly optimize its MS data acquisition, and database searching and data analysis for the TMT-labeled SCP platform using a standard cell line enzymatic mixture, which mixture are generated from HEK-293 cell lysate by filter aided sample preparation (FASP) protocol ^19^, desalted, determined concentration, and diluted to defined amount when used. To determine peptide amount in carrier channel and optimize timsTOF MS conditions, we design a set of six TMTsixplex experiments with two 50x carrier replica, two 100x carrier replica, and two 200x carrier replica. Meanwhile, each TMT EXP including one 0-cell channel (Blank channel), two single-cell-sized channels, one 2-cell-sized channel, and one 5-cell-sized channel. With the increase peptide amount of carrier channel, protein quantitative numbers in single-cell-sized, 2-cell-sized, and 5-cell-sized channels increase accordingly (Figure 3A). In each TMT EXP, the Pearson correlation coefficients of protein TMT intensities among 1:1:2:5-cell-sized channels are averagely above 0.9 (Figure 3B) with slight progressive declines in 100x carrier and 200x carrier EXPs. Compared with 200x carrier EXPs, 50x carrier and 100x carrier EXPs have much less interference of blank channels in terms of protein numbers (Figure 3A), protein TMT intensities (Figure 3C), and stabilities of defined quantitative ratios (Figure 3D). To take advantage of timsTOF MS sensitivity and avoid introducing too much interference in blank channels, we therefore choice 100x carrier to do further SCP experiments based on the above observations.

**Figure 3.**
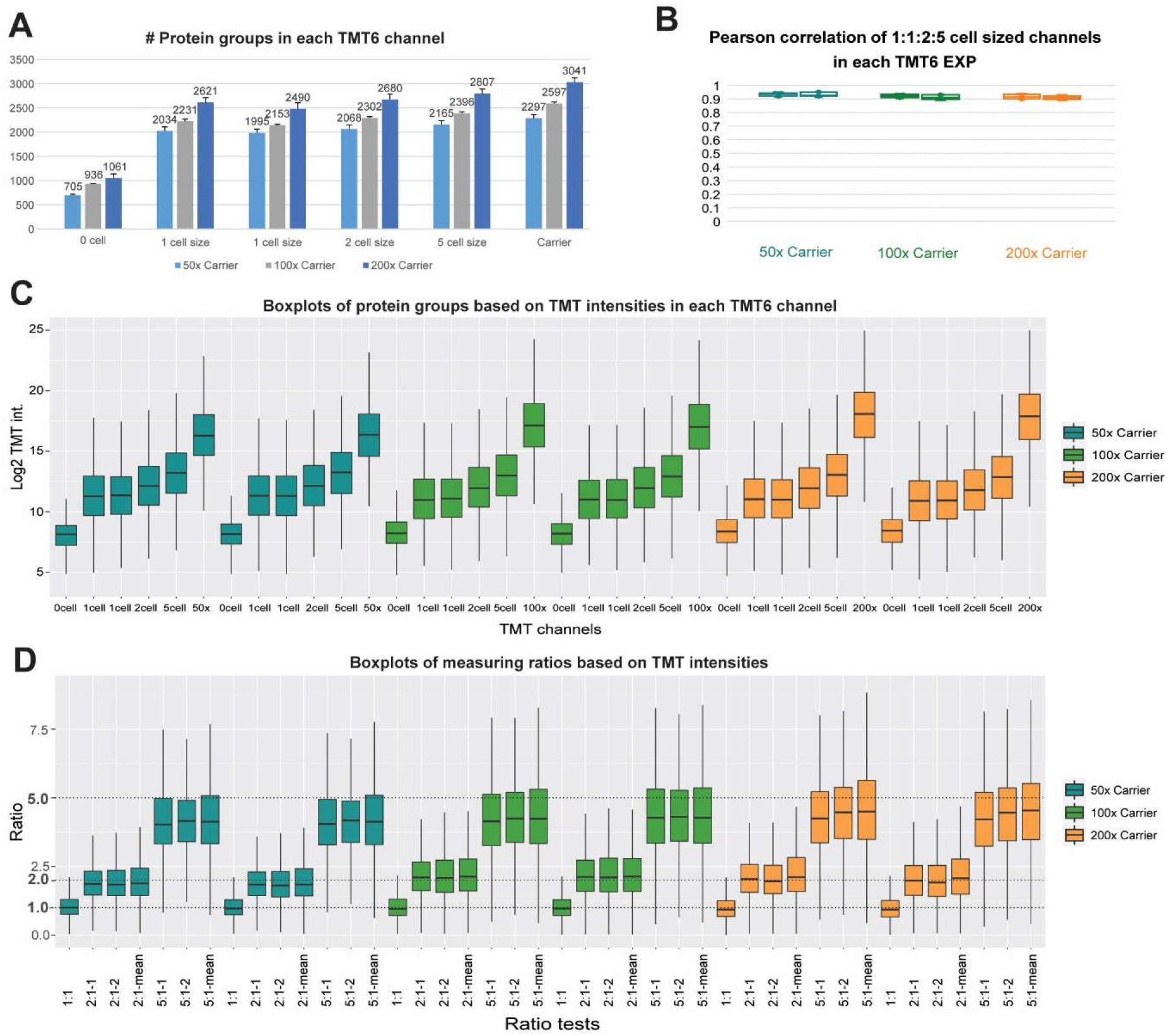
Evaluation of isobaric-labeling-based SCP pipeline using defined ratios of HEK-293T tryptic digests and timsTOF MS. With a ratio of 1:1:2:5, single-cell-sized, single-cell-sized, two-cell-sized, and two-cell-sized HEK-293T digests are labeled with four TMT channels, respectively. To optimize the most suitable carrier cell number, each TMTsixplex EXP include one empty channel and one carrier channel labeled with different amounts (50x,100x and 200x) of HEK-293T digests. As well as, different carrier TMT EXPs include a technical duplication. (A) Numbers of quantified protein groups in all TMT channels. Error bar represent standard deviations of technical replicates. (B) Pearson correlation coefficients of protein intensities are plotted for correlations between 1:1:2:5 cell-sized channels in each TMT EXP. (C) Intensity distribution of protein groups based on reporter ions across each TMT EXP in log2. (D) Boxplots of measuring ratios of protein groups based on TMT intensities between different channels in each TMT EXP. Ratio with “-mean”, the denominator is the quantitative mean of the two single-cell channels in each TMT EXP.

### Achieving small batch and large batch SCP data for Hela and HEK-293T cells

After confirmed the reliability of our UE-SCP process with QC samples (Figure 3), we expend the workflow to achieve Hela and HEK-293T SCP data. Two experimental batches have done, including a small batch (Batch1) with 8 TMT EXPs and a large batch (Batch 2) with 24 TMT EXPs. Among them, 32 or 96 single-cell TMT channels, 8 or 24 blank TMT channels, and 8 or 24 TMT carrier channels have been collected data by timsTOF MS, respectively. We summarize the results in Figure 4. The means and standard deviations of all TMT EXPs for identified protein groups, peptides, and PSMs in each batch are severally shown in Figure A-C. The average peptide-to-protein ratio among all TMT EXPs is 5.3:1 in batch 1 or 5.2:1 in batch 2, while the average identified rate (ID rate, Figure 4D) among all TMT EXPs is 26.4% in batch 1 or 25.3% in batch 2. As for single-cell TMT channels, 3220, 4003, or 4230 protein groups have been identified, while 3182, 3941, or 4165 protein groups have been quantified in batch 1 (32 cells), batch 2 (96 cells) or in total (128 cells), respectively (Figure 4E). Regarding cell groups, 3094, 3849, or 4058 protein groups have been quantified in given HEK-293T cell group, while 3135, 3883, or 4103 protein groups have been quantified in given Hela cell group (Figure 4E). As shown in Figure 4F, the quantitative orders of single-cell channels in each batch and in total have all over six orders of magnitude. Taken batch 1 and batch 2 together, Figure 4G has summarize the distribution of quantified protein group numbers in each cell group and in total. The median of quantified protein group number in 64 HEK-293T single cells, 64 Hela single cells, or all 128 single cells severally is 2222, 2306, or 2249. All these show that our SCP workflow is stable, reliable, and extremely sensitive.

**Figure 4.**
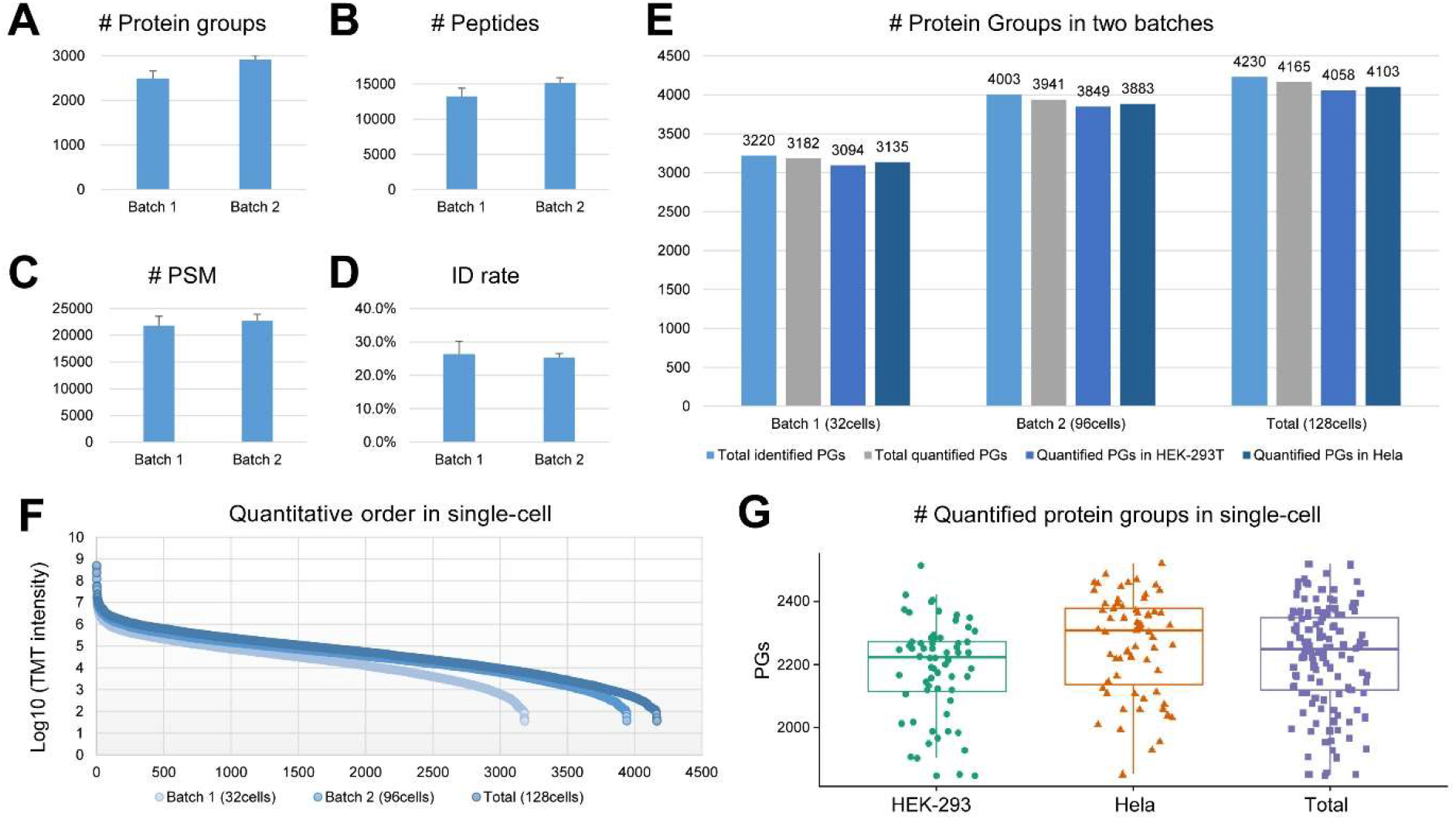
Data summary of two experimental batches. The identified protein groups (A), peptides (B), PSMs (C), ID rates (D) of all TMT EXPs in each batch are shown. ID rate, the identified rate, equals to PSM number over MS/MS number in each TMT EXPs. The identified and quantified protein group numbers of single-cell channels in each batch, each cell group, and in total are summarized (E). Quantitative orders of single-cell channels in each batch and in total are shown in panel F. Panel G has summarize the distribution of quantified protein group numbers in each cell group and in total.

### Clustering Hela and HEK-293T cell groups based on their SCP profiles

To analyze Hela or HEK-293T SCP profiles, we firstly selected the stable quantitative datasets and summarized in Figure 5A and Table S1. Among 128 single cells, the union is 3546 protein groups quantified more than 3 cells in each cell group, while the intersection is 2490 protein groups quantified more than 3 cells in each cell group and each batch. The Venn diagram (Figure 5B) has shown that the larger batch (Batch 2) reproduces almost all the quantifications of the smaller batch (Batch1), and adds more than 500 new quantified protein groups. After normalization, the 3546-protein groups SCP dataset shows good data parallelism (Figure 5C) and excellent quantitative correlation (Figure 5D). We next do unsupervised clustering for the 2490-protein groups SCP dataset. Both hierarchical cluster analysis (HCA, Figure 5E) and principal component analysis (PCA, Figure 5F) can distinct Hela and HEK-293T cell groups based on their SCP profiles. Functional annotations of the 2490 protein groups have been done using DAVID ^20^ and Metascape ^21^ bioinformatics resources. We found that extracellular exosome (GO:0070062, 37.17%) and membrane (GO:0016020, 28.24%) account for the largest and third significance in all enriched GO-CC terms, indicated outstanding cell integrities and data completeness for our SCP platform. The top 10 statistically enriched GO-CC terms were identified and the accumulative hypergeometric p-values calculated by DAVID were shown in Figure 5G and Table S2. Among the top 20 enriched functional processes (Table S3) annotated by Metascape, in addition to translation, metabolic, and energy processes, important biological pathways, such as VEGFA-VEGFR2 signaling pathway, membrane trafficking, membrane organization, and signaling by Rho GTPases, are also statistically enriched. Taken together, we strongly believe that this UE-SCP platform can obtain ultra-sensitivity, reliable quantitative, and high-throughput SCP data.

**Figure 5.**
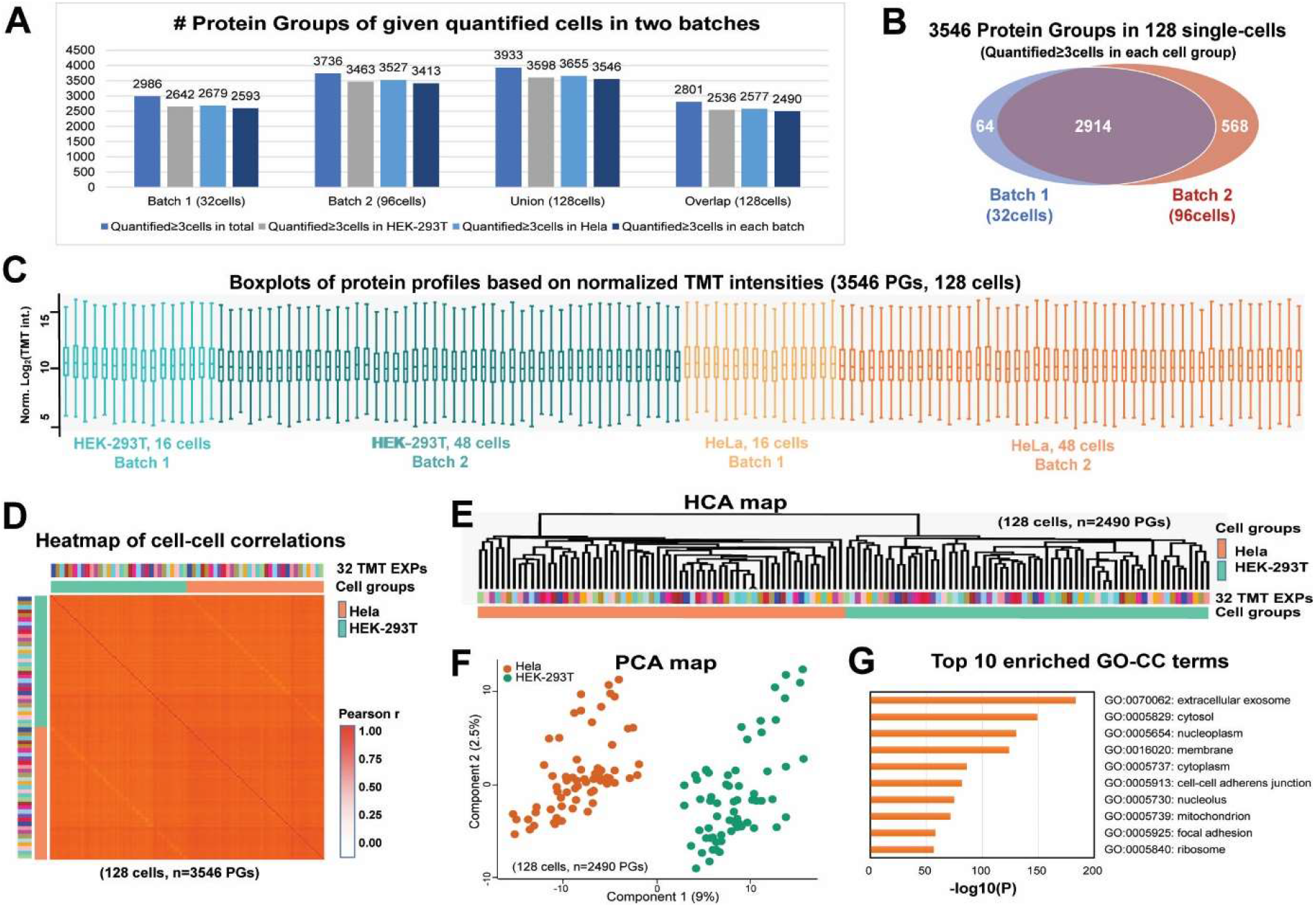
Benchmarking the quantitative SCP proteomic results in two experimental batches and overall data analysis. Reliable quantitative proteome datasets of given cells in each batch, union, and intersection were shown in panel A. Venn map (B) show the union of two batch with more than 3 cells quantified in each cell group. Normalized 3546-PGs dataset is used to draw boxplot (C) and correlation heatmap (D) for each single-cell proteome. The intersection of two batches with more than 3 cells quantified in each cell group and each batch is used to do HCA clustering (E), PCA clustering (F), and functional annotation (G).

## Discussions

In this work, we establish an ultra-sensitive and easy-to-use multiplexed single-cell proteomic pipeline. To our knowledge, it is the first application which expand stable isotope affinity labeling strategy, flexible cell sorting, and multiplexed single-cell proteomes analysis into trapped ion mobility spectrometry coupled with quadrupole time-of-flight MS. So that the numbers of quantity and quantitative protein groups are significantly improved. At the same time, we also use this pipeline in combination with Orbitrap series mass spectrometer (Data not shown). Therefore, UE-SCP can be compatible with the mainstream mass spectrometer on the market, and has universality and broad application prospects.

The UE-SCP has the following advantages: no microfluidic chip, no expensive ultrasound equipment, no difficult maintenance modifications to the liquid phase system, roboticized and accurate single-cell sorting, and quick and convenient automation. The design flux is high, which can realize the rapid collection of large-scale single-cell proteome data from single-cell sorting to data analysis. And the quantitative orders of single-cell channels are over six orders of magnitude (Figure 4F). Averagely, the identified and quantitative depth of single-cell proteome is high over 2000 protein groups per Hela or HEK-293T cell (Figure 4G). When the numbers of single cells are up to 32, 96, and 128, the depths of single-cell proteomes are over 3200, 4000, and 4200 (Figure 4E). What more, single cells from different cell lines can be clearly clustered according to their proteomic profiles (Figure 5E and 5F).

In conclusion, all of these indicate that our UE-SCP process has outstanding sensitivity, and broad application prospects in SCP research field.

## Material and Methods

### Cell culture and single-cell sorting

HEK-293T and HeLa cells were grown in DMEM (Gibco) supplemented with 10% fetal bovine serum (FBS) and 1% penicillin-streptomycin. All cells were cultured in a 37 °C incubator with 5% CO_2_. Before sorting, the adherent cells were washed twice with 1x phosphate buffered saline (PBS), detached by trypsin-EDTA (Gibco), counted, and diluted in fresh 1x PBS. Then the single HEK-293T or HeLa cell were sorted to a 384-well plate containing lysis buffer using the cellenONE® (SCIENION) equipped with the large size Piezo Dispense Capillary (PDC). Filter criteria was decided by the first 100 detected particles to exclude the broken cells, ensuring the isolated cells alive.

### Carrier peptide preparation

HEK-293T and HeLa cells were cultured as described above. Cell were harvested by SDT buffer and heated at 95 °C for 5 min to lysis. Proteins were digested with trypsin following the filter assisted sample preparation (FASP) procedure, as described previously.^19^ Briefly, 200 μg proteins from each cell line were diluted by 100 µL UA buffer (8 M urea, 150 mM Tris–HCl, pH 8.0) and loaded on an ultrafiltration filter (10 kDa cutoff, Pall Corporation). Following once wash with 100 μl UA buffer, 50 mM iodoacetamide (IAA) was added to carboxyamidomethylate thiols for 30 min in the dark. Filters were further washed twice with 100μl UA buffer and four times with 100 μl 50 mM NH_4_HCO_3_. Four μg trypsin in 100 μl 50 mM NH_4_HCO_3_ was placed on the filter to digest protein at 37 °C overnight, then another 4 μg trypsin was added to digest for 4h. Peptides were collected and the filter was wash with 100μl 50 mM NH_4_HCO_3_. The eluate was combined and desalted on a C18 homemade StageTip. Clean peptides were lyophilized and resuspended in 0.1% (v/v) formate/ddH_2_O. The concentration of the peptides was determined by the BCA assay.

### SCP proteomic sample preparation

The plate containing sorted cells was short spined and frozen at -80 °C overnight, then heated at 90 °C for 10 min to lysis cells. Protein was digested with trypsin (MS Grade, Thermo Fisher Scientific) for 4 h at 37 °C. Single-cell sample, blank sample and carrier sample were labelled with TMTsixplex™ reagents (Thermo Fisher Scientific) in 40% (v/v) ACN for 1 h at room temperature followed by quenching. The preparation process of blank sample and carrier samples were consistent with single-cell samples in the same plate. Carrier peptides were added and dispensed to the 384-well plates before single-cell sorting. Differentially labelled samples from one TMT6plex set were combined and desalted on a C18 homemade StageTip. Clean peptides were lyophilized and resuspended in 0.1% (v/v) formate/ddH_2_O for LC-MS/MS analysis.

### LC-MS/MS analysis

The LC-MS/MS analysis was performed on a trapped ion mobility spectrometry coupled to time-of-flight mass spectrometer (timsTOF MS, Bruker) combined with a high performance applied chromatographic system nanoElute® (Bruker). Peptides were loaded on to an in-house packed column (75 μm × 250 mm; 1.9 μm ReproSil-Pur C18 beads, Dr. Maisch GmbH, Ammerbuch) which was heated to 60 °C, and separated with a 60-min gradient of 2% to 80% mobile phase B at a flow rate of 300 nL/min. The mobile phases A and B were 0.1% (v/v) formate/ddH_2_O and 0.1% (v/v) formate/acetonitrile, respectively. For the analysis of isobaric labeling peptides with TMT, the mass spectrometer was operated in the parallel accumulation-serial fragmentation (PASEF) data combined with dependent acquisition (DDA) mode. The full MS scan range was set from 100 to 1700 m/z. One acquisition cycle took 100ms and contained one MS1 survey TIMS-MS and 8 PASEF MS/MS scans with two TIMS stepping.

### Database searching

DDA PASEF data were analyzed on PEAKS software (Version Online X) and searched against the UniProt/SwissProt human database (20,350 entries, May 25, 2020), which extracts features from four-dimensional isotope patterns and associated MS/MS spectra. PEAKS Online X searches were performed using standard workflow, with PEAKS DB for identification and followed by quantitative analysis using PEAKS Q. False-discovery rates were controlled at 1% on peptide and protein group levels. Peptides with a length from 6 to 45 amino acids were considered for the search including N-terminal acetylation and methionine oxidation as variable modifications and the TMTsixplex reagents as fixed modification. Enzyme specificity was set to trypsin cleaving c-terminal to arginine and lysine.

### Data analysis

The steps of aggregation, normalization, and batch effect removal of single-cell channel proteome data were as follows. The union or intersection of two batches referred to proteins quantified in more than three single cells in HEK-293T group and Hela group in both batches or in each batch. The reporter ion intensities of proteins in each single-cell channel were quantile normalized, and the missing values were filled in according to the normal distribution. Then, the ComBat function in R package SVA was used to remove batch effect among multiple TMT experiments. The obtained data matrix files were used for hierarchical cluster analysis and principal component analysis. Functional annotation based on DAVID ^20^ and Metascape ^21^ bioinformatics resources as default params.

## Acknowledgements

We thank all members of our laboratories for helpful discussions. This work was supported by the National Natural Science Foundation of China (NSFC) (82030099), the Shanghai Municipal Science and Technology Commission “Science and Technology Innovation Action Plan” technical standard project (21DZ2201700).

## Author contributions

Conceptualization, S.Y.D., L.C. and W.H.; Methodology & Investigation, L.C., G.L., L.Z.Y, S.Y.D., and W.Q.Q.; Writing – Original Draft, L.C., G.L., L.Z.Y; Writing – Review & Editing, S.Y.D., W.Q.Q., Z.H.P, and W.H.; Supervision, S.Y.D., L.C. and W.H.

